# Unravelling the effects of methylphenidate on the dopaminergic and noradrenergic functional circuits

**DOI:** 10.1101/2020.03.09.983528

**Authors:** Ottavia Dipasquale, Daniel Martins, Arjun Sethi, Mattia Veronese, Swen Hesse, Michael Rullmann, Osama Sabri, Federico Turkheimer, Neil A Harrison, Mitul A Mehta, Mara Cercignani

**Author notes:** **Corresponding Author:** Dr. Ottavia Dipasquale, Centre for Neuroimaging Sciences, Institute of Psychiatry, Psychology & Neuroscience, King’s College London, De Crespigny Park; SE5 8AF; London, UK.

## Abstract

Functional magnetic resonance imaging (fMRI) can be combined with drugs to investigate the system-level functional responses in the brain to such challenges. However, most psychoactive agents act on multiple neurotransmitters, limiting the ability of fMRI to identify functional effects related to actions on discrete pharmacological targets. We recently introduced a multimodal approach, REACT (Receptor-Enriched Analysis of functional Connectivity by Targets), which offers the opportunity to disentangle effects of drugs on different neurotransmitters and clarify the biological mechanisms driving clinical efficacy and side effects of a compound. Here, we focus on methylphenidate (MPH), which binds to the dopamine transporter (DAT) and the norepinephrine transporter (NET), to unravel its effects on dopaminergic and noradrenergic functional circuits in the healthy brain at rest. We then explored the relationship between these target-enriched resting state functional connectivity (FC) maps and inter-individual variability in behavioural responses to a reinforcement-learning task encompassing a novelty manipulation to disentangle the molecular systems underlying specific cognitive/behavioural effects.

Results showed a significant MPH-induced FC increase in sensorimotor areas in the functional circuit associated with DAT. We also found that MPH-induced variations in DAT-and NET-enriched FC were significantly correlated with inter-individual differences in effects of MPH on key behavioural responses associated with the reinforcement-learning task.

Our findings show that MPH-related FC changes are specifically associated with DAT and provide evidence that when compounds have mixed pharmacological profiles, REACT may be able to capture regional functional effects that are underpinned by the same cognitive mechanism but are related to distinct molecular targets.

## INTRODUCTION

Methylphenidate (MPH) is a psychostimulant medication widely used to treat attention deficit hyperactivity disorder (ADHD), a mental health disorder characterised by behavioural symptoms including impulsiveness, hyperactivity and inattention [1,2]. Though MPH has been used for more than half a century, we still lack a clear understanding of the exact neurochemical mechanisms through which it exerts its clinical effects [2-4].

MPH has a dual pharmacological profile inhibiting the reuptake of both dopamine (DA) and noradrenaline (NE) by blocking their respective transporters (DAT and NET respectively) [5-7]. This consequently increases the bioavailability of synaptic DA and NE [8,9]. These properties have been well documented in a rich diversity of in-vitro, animal and human studies [5]. However, despite this, it remains unclear whether the functional effects of MPH are best understood through the DAT or NET binding (for a detailed review please see [5]). Studies evaluating in-vitro activity for both these neurochemical transporters seem to agree that the racemic mixture dl-MPH typically used in clinical formulations have higher affinity and uptake inhibition activity on the DAT than the NET [5]. In-vivo human positron emission tomography (PET) studies have also shown a clear accumulation of MPH in the basal ganglia and binding to DAT (ED_50_ = 0.25 mg/kg) [10,11]. However, in the living human brain MPH has also been shown to bind to NET (ED_50_ = 0.14 mg/kg) with a higher affinity than to DAT [12,13].

A similar issue relates to effects of MPH on human cognition and brain function. Again, it is unclear whether effects mostly relate to binding to DAT, NET or both systems, and whether effects on different regional circuits are related to the engagement of these two targets, given the known differences in their distribution densities [14-18]. Several studies have combined acute administration of MPH with functional magnetic resonance imaging (fMRI) in both healthy and clinical populations to try and unravel potential effects on brain function and gain a clearer understanding of its effects on behaviour [2,3]. These studies have employed both resting state designs and task-based approaches that tap into specific cognitive constructs. Pharmacological resting state fMRI (rs-fMRI) allow the evaluation of basic pharmacodynamical effects unconstrained by the nature of any task and have been widely used [19,20]. Studies in healthy individuals have shown that acute administration of clinically relevant MPH doses induce measurable and meaningful changes in functional connectivity (FC). For example, one study found an MPH-associated increase in intrinsic connectivity between brain areas involved in sustained attention [21]. Another reported that MPH enhanced resting state FC of the striatum/thalamus with primary motor cortex and increased negative FC with frontal executive regions [22]. In another, MPH was shown to reduce the coupling within visual and somatomotor networks and increase the competitive decoupling between the default mode and task positive networks [23]. However, given that the blood-oxygen level dependent (BOLD) signal underpinning fMRI studies has no intrinsic selectivity to any particular neurochemical target [24,25], it is still unclear which MPH pharmacological targets underpin observed changes in brain FC.

To bridge this gap, we have previously developed a novel multimodal method (Receptor-Enriched Analysis of functional Connectivity by Targets - REACT) which enriches rs-fMRI analyses with information about the distribution density of molecular targets derived from PET imaging. Further, we have shown that the functional effects of 3,4-Methyl-enedioxy-methamphetamine (MDMA) can be understood through the distribution of its main serotonergic targets [26]. Here, we applied REACT to multi-echo rs-fMRI data acquired in a cohort of healthy participants after an acute challenge of MPH to investigate how drug-related changes in resting state FC relate to the distribution of the DAT and NET. Given the considerable affinity of MPH to both neurotransmitters [2], we hypothesised that FC informed by these two targets would be sensitive to MPH effects.

We also performed an exploratory analysis to test whether MPH-induced changes in the DAT- and NET-enriched FC can be linked to inter-individual differences in behavioural responses on a reinforcement-learning task with known sensitivity to MPH (results for the main effects of this task are reported elsewhere [27]). Multiple fMRI studies have shown that inter-individual differences in behavioural responses to a task can be related to individual differences in spontaneous cortical activity at rest [28,29]. Our hypothesis is that by using the functional maps related to the MPH targets, we can link inter-individual variability in behavioural response to the task with the resting state FC and tease apart the variability associated with DAT and NET.

## METHODS

### Participants and study design

The dataset used in this work is part of a larger study on ADHD [27]. We included data from thirty healthy controls (HC, 33 ± 9.5 years, M/F: 19/11). Participants were recruited using on-line classified advertising websites and university mailing lists. Local and national ethical approvals were obtained from Brighton and Sussex Medical School (14/014/HAR; 12/131/HAR) and the East of England (Hertfordshire) National Research Ethics Committee (reference: 12/EE/0256). All participants provided written informed consent.

We followed a randomized, within-subjects, double-blind, placebo-controlled design where participants received 20 mg of oral MPH or placebo in two separate sessions, spaced by a minimum of 1 week. This dose of MPH is at the lower end of the clinical dose and allow us to evaluate functional effects in the brain within a clinically relevant dose while keeping potential side effects at minimum.

We excluded one subject because of excessive head movement during the placebo session.

### Image acquisition

Ninety minutes after drug dosing, participants completed an MRI session, to coincide with peak effects of MPH on DAT occupancy [11]. The session included a 9-minute multi-echo rs-fMRI scan. The rs-fMRI scan was used for the analysis presented herein. Prior to the resting state scan were three runs of a reinforcement-learning task encompassing a novelty manipulation [30,31], which is reported in detail elsewhere [27]. For the purpose of this work we use the behavioural outcome of the task independent of the fMRI data concurrently acquired.

Data were acquired using a 1.5 T Siemens Avanto MRI scanner (Siemens AG Medical Solutions, Erlangen, Germany) equipped with a 32-channel head-coil. Rs-fMRI data were obtained using a T2*-weighted multi-echo EPI sequence [32] (TR = 2570 ms; TEs = 15, 34, 54 ms; flip angle = 90°; resolution = 3.7 × 3.75, slice thickness = 4.49 mm; matrix size = 64 × 64; 31 axial slices; 200 volumes). A 3D T1-weighted anatomical scan was obtained for each participant in one session using an MP-RAGE acquisition (TR = 2730 ms, TE = 3.57 ms, TI = 1000 ms, flip angle = 7°, matrix = 256 × 240, number of partitions = 192, GRAPPA factor = 2, resolution = 1 mm^3^).

### Image pre-processing

The rs-fMRI dataset was pre-processed using AFNI [33] and FMRIB Software Library (FSL). Pre-processing steps included volume re-alignment, time-series de-spiking and slice time correction. After the pre-processing, functional data were optimally combined (OC) by taking a weighted summation of the three echoes using an exponential T2* weighting approach [34]. The OC data were then de-noised with the Multi-Echo ICA (ME-ICA) approach implemented by the tool meica.py (Version v2.5 beta9) [35,36], given its proven effectiveness in reducing non-BOLD artefacts and increasing the temporal signal-to-noise ratio [36-38]. White matter and cerebrospinal fluid signals were regressed out and a high-pass temporal filter with a cut-off frequency of 0.005 Hz was applied.

A study-specific template representing the average T1-weighted anatomical image across subjects was built using the Advanced Normalization Tools (ANTs) [39]. Each participant’s dataset was co-registered to its corresponding structural scan, then normalized to the study-specific template before warping to standard MNI152 space. Images were finally resampled at 2×2×2 mm^3^ resolution.

### Population-based molecular templates

For the analysis with REACT, we used molecular templates of the DAT and NET systems. The DAT map is a publicly available template of ^123^I-Ioflupane SPECT images (https://www.nitrc.org/projects/spmtemplates) from 30 HC without evidence of nigrostriatal degeneration [40]. The NET atlas was obtained by averaging the [^11^C]MRB PET brain parametric maps from an independent dataset of 10 HC (33.3 +/-10 years, four women). More details on the processing of the PET data can be found in [41]. These two atlases were already standardised to MNI space with 2 mm^3^ resolution. They were further normalised to scale the image values between 0 and 1, although preserving the original intensity distribution of the images, and masked using a standard grey matter (GM) mask. This mask comes from a probabilistic GM map available in FSL, which we thresholded in order to keep only the voxels with >30% probability of being GM, and binarized. Of note, occipital areas (defined using the Harvard Oxford Atlas) were masked out for both molecular atlases as they were used as reference regions for quantification of the molecular data in the kinetic models for the radioligands.

### Functional connectivity analysis with the REACT method

The functional circuits related to the DAT and the NET systems were estimated with REACT using a two-step multivariate regression analysis [42,43] implemented in FSL (fsl_glm command). This analysis is conceptually comparable to a dual-regression, often used in rs-fMRI studies to investigate the FC of the resting state networks. The main difference with this standard approach is that, with REACT, molecular templates are used in place of the resting state networks as a set of spatial regressors in the first multivariate regression analysis. The DAT and NET templates are entered into the first step of this analysis to calculate the dominant BOLD fluctuation within these maps [44]. Both rs-fMRI data and the design matrix were demeaned (--demean option). The rs-fMRI volumes were masked using a binarized atlas derived from the molecular data to restrict the analysis to the voxels for which the transporter density information was available in the template. The subject-specific time series estimated in this first step were then used as temporal regressors in a second multivariate regression analysis to estimate the subject-specific spatial maps of the BOLD response after MPH and placebo. At this stage, the analysis was conducted on the whole grey matter volume. Both data and the design matrix were demeaned (--demean option); the design matrix columns were also normalised to unit standard deviation with the --des_norm option [42]. The general framework of this analysis has been reported elsewhere [26].

### Statistical analysis

In order to test our main hypothesis, we run a paired t-test to compare the subject-specific target-enriched spatial maps of the two drug conditions (MPH and placebo). We applied cluster-based inference within Randomise [45], using 5000 permutations per test and contrast. A cluster was considered significant if p_FWE_ < 0.05, corrected for multiple comparisons using the threshold-free cluster enhancement (TFCE) option [46], Bonferroni-corrected for multiple comparisons across maps (DAT and NET) and contrasts (MPH > placebo, placebo > MPH).

We also conducted some further post-hoc analyses to explore whether there were any areas of the brain where our DAT and NET-enriched FC maps were related to some of the key-behavioural responses elicited during the task the subjects performed immediately before the resting state scan. For that, we conducted a set of linear regression analyses between the target-enriched FC maps (MPH minus placebo) and four behavioural scores: 1) Overall task performance (£ won); 2) reward-learning rate (higher score reflects faster updating of reward values with experience); 3) persistence in selecting novel options after their first appearance; 4) persistence in selecting novel rather than familiar stimuli after their first appearance. For each regression analysis, we used the difference between scores on active drug and placebo (MPH minus Placebo). These tests were also performed with Randomise using 5000 permutations per test and contrast. For the significant correlations, we then extracted the mean FC value (MPH minus Placebo) from the clusters we found to be significantly associated with the behavioural responses in order to estimate the Pearson correlation coefficient using SPSS. Please note that these values are simply presented as a measure of the effect size. We did not conduct statistical inference as such analysis would be circular.

## RESULTS

In Figure 1, we show the templates of the molecular density distribution of the DAT (left panel, top row) and NET maps (right panel, top row) that we used in the REACT analysis. We also show the weighted-maps of their respective functional-associated circuits for each drug condition, averaged across participants.

**Fig. 1.**
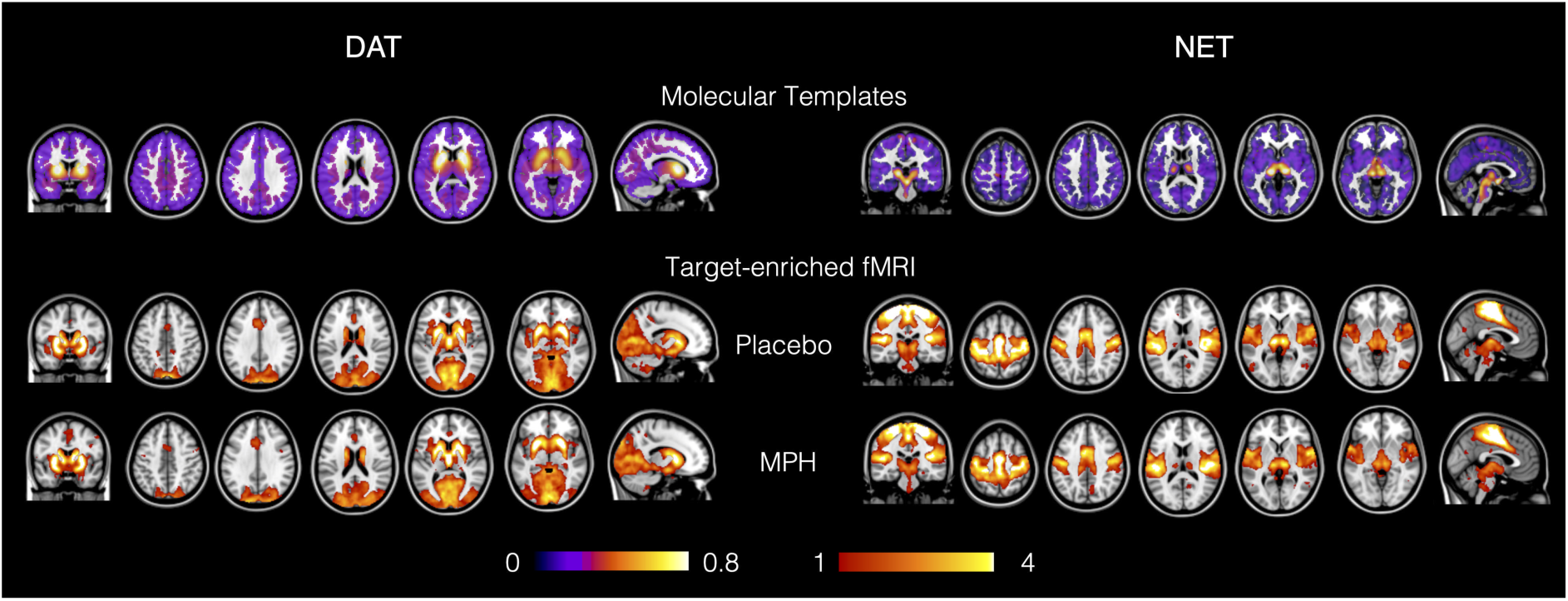
Maps of the molecular templates of the dopamine and noradrenaline transporters (DAT and NET) and their respective target-enriched fMRI maps. The fMRI maps are averaged across subjects for the placebo and methylphenidate (MPH) conditions. The occipital areas were masked out for both molecular atlases since they were used as reference regions for the quantification of the molecular data in the kinetic models for the radioligands.

We found a significant increase in FC (p_FWE_ < 0.05, corrected for multiple comparisons at the cluster level using the threshold-free cluster enhancement (TFCE) option, Bonferroni-corrected for multiple comparisons across maps and contrasts) after MPH in the maps enriched by the DAT distribution (Fig. 2). Specifically, this effect mainly involved sensorimotor areas including the precentral and postcentral gyri and the anterior division of the supramarginal gyrus. No changes between MPH and placebo were found in the NET-enriched FC maps.

**Fig. 2.**
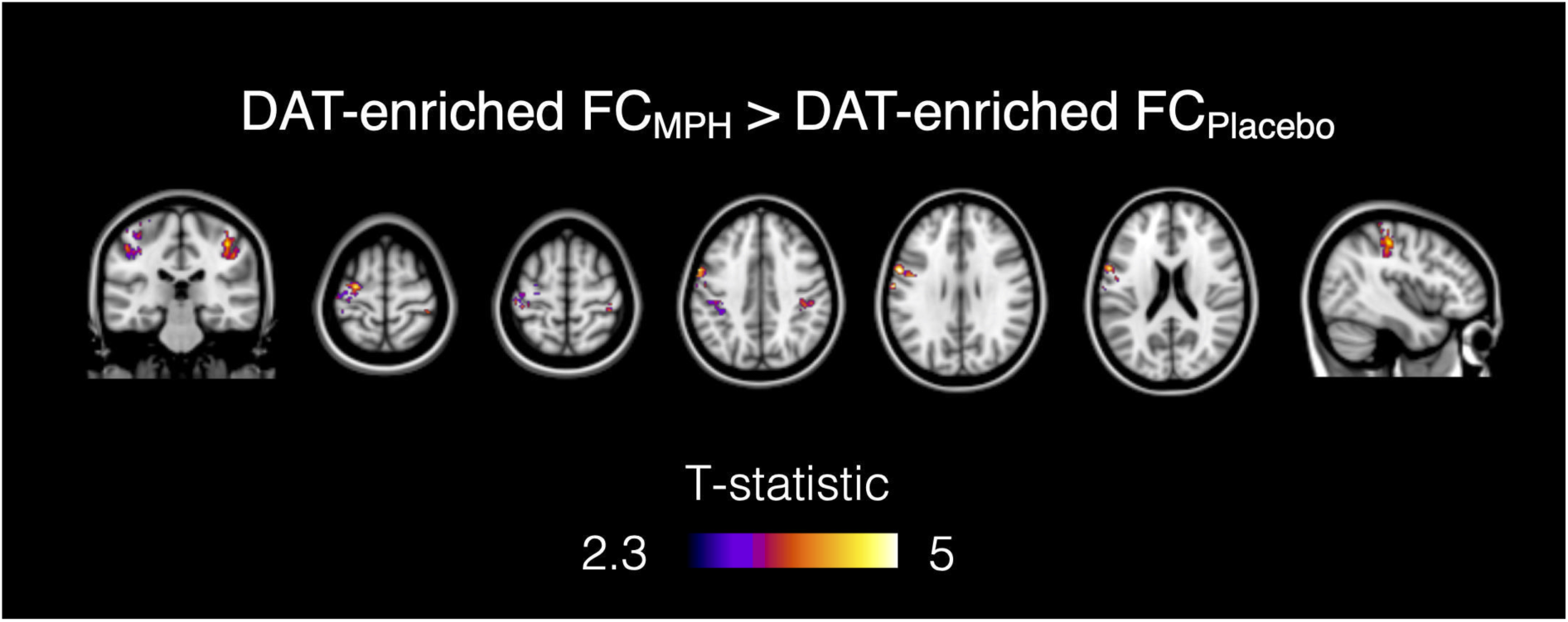
Functional connectivity (FC) changes after MPH compared to placebo in the DAT-enriched maps. The MPH-induced FC increase is localised in the precentral and postcentral gyri and the anterior division of the supramarginal gyrus.

### Relationship between drug-induced functional changes at rest and behavioural responses during the novelty reinforcement learning task

We found significant negative correlations between persistence in selecting novel options after their first appearance (MPH minus placebo) and the MPH-induced functional changes in the DAT and the NET-enriched FC maps (MPH minus placebo). Specifically, we found a negative correlation between DAT-enriched FC and this behavioural score in the cerebellum, i.e. the right crus I and II, the right VI and VIIb and the vermis crus II (r = −0.739, 95% CI = −0.865 to −0.542; Fig. 3A). For the NET-enriched FC, this correlation mainly involved the precentral gyrus, the posterior cingulate cortex and precuneus (r = −0.687, 95% CI = −0.831 to −0.488; Fig. 3A). We also found a positive association between FC changes induced by MPH in the NET-enriched maps and the task performance (£ won) in the lateral occipital cortex, precuneus and superior parietal lobule (r = 0.753, 95% CI = 0.590 to 0.857; Fig. 3B). We did not find any other significant correlation for any of the remaining behavioural measures tested.

**Fig. 3.**
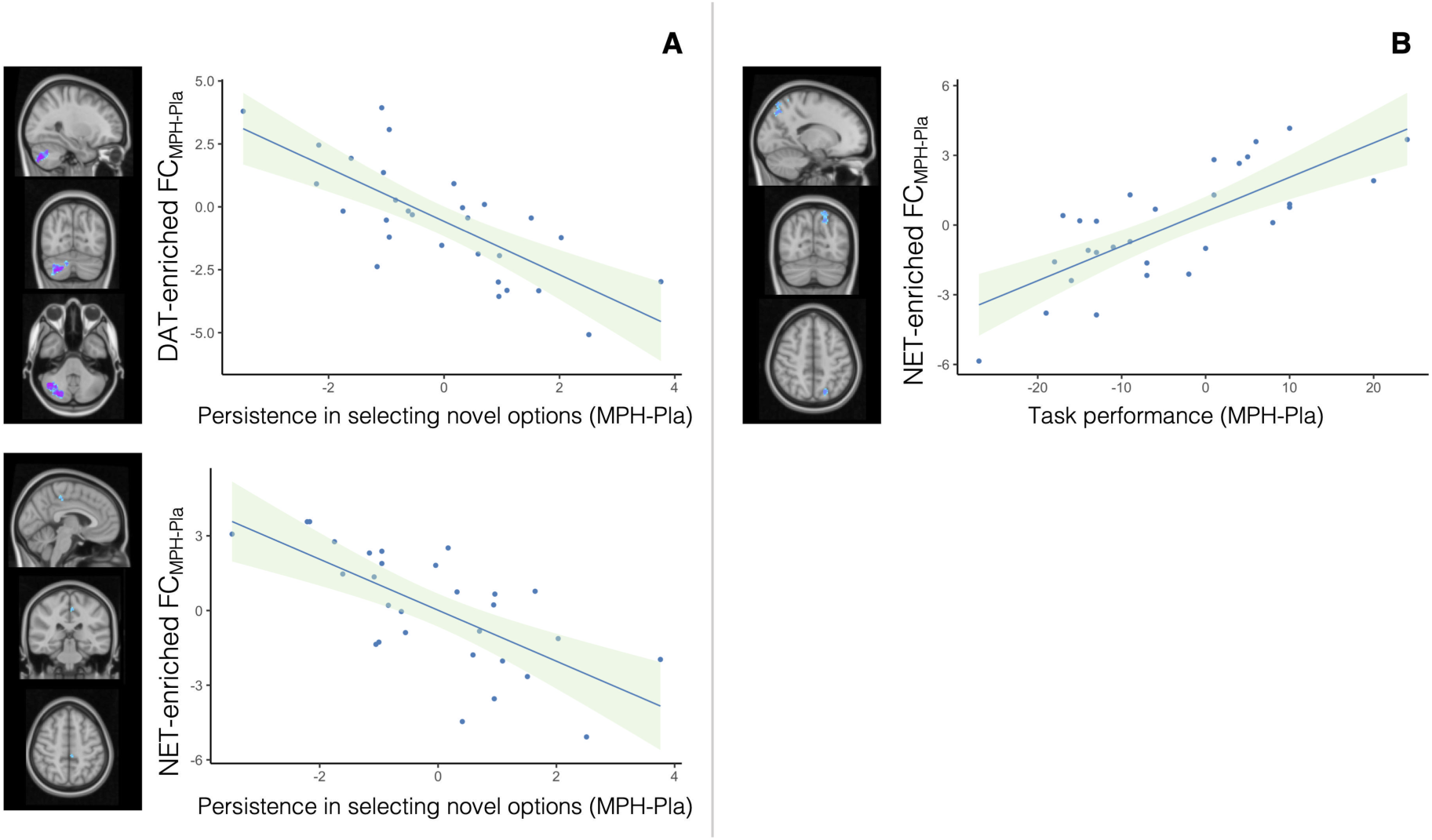
Correlations between behavioural response and functional connectivity (FC). Panel A: The persistence in selecting novel rather than familiar stimuli after their first appearance (MPH minus placebo) is inversely correlated with FC changes (MPH minus placebo) in the DAT-enriched and NET-enriched FC maps. Panel B: the task performance (MPH minus placebo) is positively correlated with functional increases in the NET-enriched FC maps.

## DISCUSSION

In this study, we applied our recently developed multimodal method for FC analysis informed by molecular targets (REACT) to investigate how drug-related changes in resting FC after a single administration of MPH in healthy individuals relate to the distribution of its main targets, i.e. the DAT and NET. In line with our main hypothesis, we found that MPH changed connectivity within the functional network related to the DAT. However, we did not find any significant drug-effects related to the NET distribution at the robust statistical thresholds reported here. Furthermore, we found that different regional changes in the DAT and NET-enriched FC maps can significantly explain some of the inter-individual variance in participants’ behaviour during a novelty reinforcement learning task engaging both systems [47]. Moreover, we have extended our method validation pipeline by showing that REACT is able to capture meaningful changes in the target-enriched functional circuits beyond those related to the serotonin system, which we reported in our first proof-of-concept work [26].

Here we were able to show that MPH effects on resting FC co-vary with the known distribution of one of its main targets, the DAT. We found FC increases within the DAT-related network mapping somatomotor areas, such as the precentral and postcentral gyri and the anterior division of the supramarginal gyrus. This overlaps with regions where regional cerebral blood flow measured at rest is modulated by a single dose of MPH [48] as part of a pattern which also includes the caudate nucleus, thalamus and mid-brain, all areas within the DAT-enriched connectivity map. Our findings are also consistent with previous observations in healthy individuals that MPH increases the FC between the striatum (a brain region highly enriched in DAT) and the somatomotor cortex [22]. Our results also mirror previous studies where levodopa and haloperidol (pro- and anti-dopaminergic drugs, respectively) were found to modulate the resting state and task-related FC between the motor cortex and the striatum in healthy participants [49,50]. While our data was acquired at rest, and therefore making inferences about the potential behavioural implications of such findings is inherently speculative, we note that the FC increase in the somatomotor cortex is in line with some reported effects of MPH and other catecholaminergic agents on motor performance [51]. For instance, MPH has been shown to improve motor functions in clinical conditions encompassing motor deficits. In fact, a single dose of MPH has been shown to improve motor coordination in children with developmental coordination disorder and ADHD [52], and low doses of this drug have been reported to improve gait and voluntary movement in patients with Parkinson disease [53].

The lack of findings for the NET-enriched functional circuit is surprising for many reasons. First, human PET studies have shown that at clinical doses (such as those used herein), MPH binds to the NET with a higher affinity than to the DAT [12,13]. Second, there is further evidence from studies in experimental animal showing that MPH can directly affect the discharge properties of the locus coeruleus (LC) [54], which is the main source of noradrenergic innervation in the brain; these findings are supported by a rs-fMRI study in humans showing that MPH exerts effects on the FC of the LC [55]. Third, there is a general consensus that the MPH pro-cognitive effects in ADHD patients and HC involve modulation of the prefrontal cortex (PFC) [56]. While both DA and NA have critical influence on PFC cognitive functioning [56,57], there are relatively low levels of DAT in the PFC [58], supporting the hypothesis that MPH and other psychomotor stimulants effects in this area may involve the NET inhibition [59]. However, a number of reasons can be advanced to explain this lack of findings in the NET-related FC. First, there may be a limitation in our technique. We use the available binding sites for NET, but PET occupancy studies suggest a regional variation in the MPH occupancy of NET [12] which would weigh more towards the effects at the LC which may be poorly represented in the MRI maps acquired at 1.5 or 3T [60]. A general limitation that we should also consider is that our subjects performed a reinforcement learning task – which is known to elicit a dopaminergic response [61] – immediately before the rs-fMRI scan, and that this may have induced some carry-over effects biasing FC towards the DAT-related network. However, it is important to point out that the reuptake of dopamine and noradrenaline is complex, as they can both participate in the transport of DA and NA [62,63]. In areas of low DAT density, the affinity of dopamine to NET can be even higher than its affinity to DAT itself [64,65]. This implies that MPH could still affect the noradrenergic transmission through DAT inhibition. Moreover, we cannot exclude that some NET-related FC changes that do not reach statistical significance at rest may emerge during a paradigm preferentially engaging attentional networks [66,67]. Finally, we acquired data on a single time-interval post-dosing with a single low dose. PET occupancy of 10mg and 40mg MPH at NET assessed in vivo using [^11^C]MRB indicates that this dose is sufficient to produce a significant NET occupancy with robust peak effects between 75 minutes and at least 3 hours post-dosing [13].

Although MPH did not elicit robust behavioural responses during the novelty-reinforcement learning task in HC (data published elsewhere [27]), we found that differences between MPH and placebo on DAT- and NET-enriched FC are significantly correlated with inter-individual differences in the MPH-induced changes in key-behavioural responses associated with this task. Indeed, our findings show that despite the relatively low density of DAT in the cerebellum, the DAT-enriched connectivity in this same area was related to performance. This may be related to the strong anatomical connectivity between the cerebellum and the DAT-enriched basal ganglia [68], or it may relate to direct effects within the cerebellum. Despite the low detection of DAT availability in vivo in humans, DAT is present in the cerebellum [69] and our findings match the current models suggesting a role of the cerebellum in error/novelty detection [70-72].

While we did not find a treatment effect of MPH in the NET-related FC, we did find a positive association between FC changes induced by MPH in the NET-enriched maps and overall task performance (£ won) in the lateral occipital cortex, precuneus and superior parietal lobule. We also found negative correlations between persistence in selecting novel rather than familiar options and NET-enriched FC changes induced by MPH mainly in the precentral gyrus, the posterior cingulate cortex and the precuneus. Consistent with our findings, these areas have previously been shown to be implicated in novelty processing [73-75]. Animal studies have shown that the occipital and parietal cortices receive a dense noradrenergic innervation and that pharmacological NET blockade increases extracellular NE (but not DA) in these areas [76,77]. Furthermore, a resting state fMRI study in healthy subjects reported a FC decrease induced by atomoxetine, i.e. a relatively selective NET blocker, predominantly in the posterior brain regions including the visual system [78]. Another fMRI study using a n-back task in healthy volunteers reported atomoxetine-induced FC changes in the frontoparietal network, including areas such as the precentral gyrus and the precuneus, during working-memory processing [79].

Although preliminary, our findings suggest that REACT may hold the potential to tease apart regional functional effects related to different molecular targets underlying the same cognitive/behavioural downstream effect when compounds like MPH binding with considerable affinity to more than one molecular target are used. However, since we relied on correlations between resting state FC and behavioural measures acquired at a different time point, this hypothesis would benefit from further development of REACT to accommodate task-based designs.

Our study has some limitations to acknowledge. First, we did not collect blood samples and assay plasma levels of MPH to control for individual differences in drug exposure or to explore pharmacokinetic/pharmacodynamic relationships. Second, while the original study also included patients with ADHD [27], we only report data from the healthy control group, where all participants received the same drug at the same dose; therefore, the implications of our findings should not be generalized to clinical populations. However, we decided to not include the ADHD dataset because all patients were on chronic stimulant treatment, which is known to produce long-lasting changes in the baseline DAT bioavailability [80,81]. Furthermore, ADHD patients were tested under their routine treatment, which was heterogeneous (i.e. either MPH or atomoxetine). These two factors would have made the interpretation of potential findings challenging. Third, stimulants can potentially influence fMRI BOLD signal, which depends on the haemodynamic coupling of neuronal activities and local changes in blood flow and oxygenation [24]. However, previous work has found that while stimulants can decrease cortical blood flow, they do not obscure BOLD signals or disrupt neurovascular coupling during resting brain activity [82]. Finally, we relied on group-based molecular templates estimated in two independent cohorts of healthy individuals. Therefore, further specification from intra-regional variation across subjects is not possible using the current dataset as it would require PET data for each ligand and participant. Nevertheless, it would be interesting to test whether subject-specific receptor density maps would provide additional information. This last aspect may be critical moving forward to examine MPH-induced FC changes in clinical populations (i.e. ADHD), who may have alterations in the distribution of DAT [81] and NET [83].

## CONCLUSION

Using our recently developed multimodal method for FC analysis informed by molecular targets (REACT), we show that meaningful MPH effects on human brain resting FC can be understood through the distribution of, at least, one of its main targets, i.e. DAT. We also provide evidence to support the idea that this method may be able to capture concomitant differential regional functional effects related to different targets underlying the same cognitive/behavioural effect, when compounds have mixed pharmacological profiles. By defining the target-specific topography of the functional effects of pharmacological compounds in the human brain, our method holds the potential to advance our understanding of the mechanisms of action of many drugs for which target affinity is relatively well characterized but system-level brain pharmacodynamic models related to their targets are missing.

## REFERENCES

1 Storebo OJ, Pedersen N, Ramstad E, Kielsholm ML, Nielsen SS, Krogh HB, et al. Methylphenidate for attention deficit hyperactivity disorder (ADHD) in children and adolescents-assessment of adverse events in non-randomised studies. Cochrane Database of Systematic Reviews. 2018(5).

2 Faraone SV. The pharmacology of amphetamine and methylphenidate: Relevance to the neurobiology of attention-deficit/hyperactivity disorder and other psychiatric comorbidities. Neuroscience and Biobehavioral Reviews. 2018;87:255–70.

3 Czerniak SM, Sikoglu EM, King JA, Kennedy DN, Mick E, Frazier J, et al. Areas of the Brain Modulated by Single-Dose Methylphenidate Treatment in Youth with ADHD During Task-Based fMRI: A Systematic Review. Harvard Review of Psychiatry. 2013;21(3):151–62.

4 Grunblatt E, Bartl J, Schmid R, Walitza S. Methlyphenidate treatment in attention-deficit hyperactivity disorder: What do we know about the mechanism of action of methylphenidate? Eur Child Adoles Psy. 2013;22:S188–S88.

5 Markowitz JS, Patrick KS. Differential pharmacokinetics and pharmacodynamics of methylphenidate enantiomers: does chirality matter? J Clin Psychopharmacol. 2008;28(3 Suppl 2):S54–61.

6 Williard RL, Middaugh LD, Zhu HJ, Patrick KS. Methylphenidate and its ethanol transesterification metabolite ethylphenidate: brain disposition, monoamine transporters and motor activity. Behav Pharmacol. 2007;18(1):39–51.

7 Markowitz JS, DeVane CL, Pestreich LK, Patrick KS, Muniz R. A comprehensive in vitro screening of d-, l-, and dl-threo-methylphenidate: an exploratory study. J Child Adolesc Psychopharmacol. 2006;16(6):687–98.

8 Koda K, Ago Y, Cong Y, Kita Y, Takuma K, Matsuda T. Effects of acute and chronic administration of atomoxetine and methylphenidate on extracellular levels of noradrenaline, dopamine and serotonin in the prefrontal cortex and striatum of mice. J Neurochem. 2010;114(1):259–70.

9 Rowley HL, Kulkarni RS, Gosden J, Brammer RJ, Hackett D, Heal DJ. Differences in the neurochemical and behavioural profiles of lisdexamfetamine methylphenidate and modafinil revealed by simultaneous dual-probe microdialysis and locomotor activity measurements in freely-moving rats. Journal of Psychopharmacology. 2014;28(3):254–69.

10 Volkow ND, Fowler JS, Wang G, Ding Y, Gatley SJ. Mechanism of action of methylphenidate: insights from PET imaging studies. J Atten Disord. 2002;6 Suppl 1: S31–43.

11 Volkow ND, Wang GJ, Fowler JS, Gatley SJ, Logan J, Ding YS, et al. Dopamine transporter occupancies in the human brain induced by therapeutic doses of oral methylphenidate. Am J Psychiatry. 1998;155(10):1325–31.

12 Hannestad J, Gallezot JD, Planeta-Wilson B, Lin SF, Williams WA, van Dyck CH, et al. Clinically Relevant Doses of Methylphenidate Significantly Occupy Norepinephrine Transporters in Humans In Vivo. Biological Psychiatry. 2010;68(9):854–60.

13 Hannestad J, Planeta-Wilson B, Gallezot J, Lin S, Williams W, van Dyck C, et al. Duration and degree of norepinephrine transporter occupancy by oral methylphenidate with (S,S)-[11C]MRB in healthy subjects. Journal of Nuclear Medicine. 2009;50.

14 Ciliax BJ, Drash GW, Staley JK, Haber S, Mobley CJ, Miller GW, et al. Immunocytochemical localization of the dopamine transporter in human brain. The Journal of comparative neurology. 1999;409(1):38–56.

15 Hall H, Halldin C, Guilloteau D, Chalon S, Emond P, Besnard J, et al. Visualization of the dopamine transporter in the human brain postmortem with the new selective ligand [125I]PE2I. Neuroimage. 1999;9(1):108–16.

16 Volkow ND, Ding YS, Fowler JS, Wang GJ, Logan J, Gatley SJ, et al. A new PET ligand for the dopamine transporter: studies in the human brain. J Nucl Med. 1995;36(12):2162–8.

17 Smith HR, Beveridge TJ, Porrino LJ. Distribution of norepinephrine transporters in the non-human primate brain. Neuroscience. 2006;138(2):703–14.

18 Arakawa R, Okumura M, Ito H, Seki C, Takahashi H, Takano H, et al. Quantitative analysis of norepinephrine transporter in the human brain using PET with (S,S)-18F-FMeNER-D2. J Nucl Med. 2008;49(8):1270–6.

19 Wandschneider B, Koepp MJ. Pharmaco fMRI: Determining the functional anatomy of the effects of medication. Neuroimage-Clin. 2016;12:691–97.

20 Smitha KA, Akhil Raja K, Arun KM, Rajesh PG, Thomas B, Kapilamoorthy TR, et al. Resting state fMRI: A review on methods in resting state connectivity analysis and resting state networks. Neuroradiol J. 2017;30(4):305–17.

21 Mueller S, Costa A, Keeser D, Pogarell O, Berman A, Coates U, et al. The Effects of Methylphenidate on Whole Brain Intrinsic Functional Connectivity. Human Brain Mapping. 2014;35(11):5379–88.

22 Farr OM, Zhang S, Hu S, Matuskey D, Abdelghany O, Malison RT, et al. The effects of methylphenidate on resting-state striatal, thalamic and global functional connectivity in healthy adults. International Journal of Neuropsychopharmacology. 2014;17(8):1177–91.

23 Sripada CS, Kessler D, Welsh R, Angstadt M, Liberzon I, Phan KL, et al. Distributed effects of methylphenidate on the network structure of the resting brain: A connectomic pattern classification analysis. Neuroimage. 2013;81:213–21.

24 Attwell D, Iadecola C. The neural basis of functional brain imaging signals. Trends in Neurosciences. 2002;25(12):621–25.

25 Khalili-Mahani N, Rombouts SARB, van Osch MJP, Duff EP, Carbonell F, Nickerson LD, et al. Biomarkers, Designs, and Interpretations of Resting-State fMRI in Translational Pharmacological Research: A Review of State-of-the-Art, Challenges, and Opportunities for Studying Brain Chemistry. Human Brain Mapping. 2017;38(4):2276–325.

26 Dipasquale O, Selvaggi P, Veronese M, Gabay AS, Turkheimer F, Mehta MA. Receptor-Enriched Analysis of functional connectivity by targets (REACT): A novel, multimodal analytical approach informed by PET to study the pharmacodynamic response of the brain under MDMA. Neuroimage. 2019;195:252–60.

27 Sethi A, Voon V, Critchley HD, Cercignani M, Harrison NA. A neurocomputational account of reward and novelty processing and effects of psychostimulants in attention deficit hyperactivity disorder. Brain. 2018;141(5):1545–57.

28 Baldassarre A, Lewis CM, Committeri G, Snyder AZ, Romani GL, Corbetta M. Individual variability in functional connectivity predicts performance of a perceptual task. Proc Natl Acad Sci U S A. 2012;109(9):3516–21.

29 Hampson M, Driesen NR, Skudlarski P, Gore JC, Constable RT. Brain connectivity related to working memory performance. J Neurosci. 2006;26(51):13338–43.

30 Djamshidian A, O’Sullivan SS, Wittmann BC, Lees AJ, Averbeck BB. Novelty seeking behaviour in Parkinson’s disease. Neuropsychologia. 2011;49(9):2483–8.

31 Wittmann BC, Daw ND, Seymour B, Dolan RJ. Striatal activity underlies novelty-based choice in humans. Neuron. 2008;58(6):967–73.

32 Poser BA, Versluis MJ, Hoogduin JM, Norris DG. BOLD contrast sensitivity enhancement and artifact reduction with multiecho EPI: parallel-acquired inhomogeneity-desensitized fMRI. Magn Reson Med. 2006;55(6):1227–35.

33 Cox RW. AFNI: software for analysis and visualization of functional magnetic resonance neuroimages. Computers and Biomedical research. 1996;29(3):162–73.

34 Posse S, Wiese S, Gembris D, Mathiak K, Kessler C, Grosse-Ruyken ML, et al. Enhancement of BOLD-contrast sensitivity by single-shot multi-echo functional MR imaging. Magnetic Resonance in Medicine. 1999;42(1):87–97.

35 Kundu P, Santin MD, Bandettini PA, Bullmore ET, Petiet A. Differentiating BOLD and non-BOLD signals in fMRI time series from anesthetized rats using multi-echo EPI at 11.7 T. NeuroImage. 2014;102:861–74.

36 Kundu P, Brenowitz ND, Voon V, Worbe Y, Vertes PE, Inati SJ, et al. Integrated strategy for improving functional connectivity mapping using multiecho fMRI. Proceedings of the National Academy of Sciences of the United States of America. 2013;110(40):16187–92.

37 Dipasquale O, Sethi A, Lagana MM, Baglio F, Baselli G, Kundu P, et al. Comparing resting state fMRI de-noising approaches using multi- and single-echo acquisitions. Plos One. 2017;12(3):e0173289.

38 Kundu P, Benson BE, Baldwin KL, Rosen D, Luh WM, Bandettini PA, et al. Robust resting state fMRI processing for studies on typical brain development based on multi-echo EPI acquisition. Brain Imaging Behav. 2015;9(1):56–73.

39 Avants BB, Tustison NJ, Song G, Cook PA, Klein A, Gee JC. A reproducible evaluation of ANTs similarity metric performance in brain image registration. Neuroimage. 2011;54(3):2033–44.

40 García-Gómez FJ, García-Solís D, Luis-Simón FJ, Marín-Oyaga VA, Carrillo F, Mir P, et al. [Elaboration of the SPM template for the standardization of SPECT images with 123I-Ioflupane]. Rev Esp Med Nucl Imagen Mol. 2013;32(6):350–6.

41 Hesse S, Becker GA, Rullmann M, Bresch A, Luthardt J, Hankir MK, et al. Central noradrenaline transporter availability in highly obese, non-depressed individuals. Eur J Nucl Med Mol Imaging. 2017;44(6):1056–64.

42 Filippini N, MacIntosh BJ, Hough MG, Goodwin GM, Frisoni GB, Smith SM, et al. Distinct patterns of brain activity in young carriers of the APOE-epsilon4 allele. Proc Natl Acad Sci U S A. 2009;106(17):7209–14.

43 Nickerson LD, Smith SM, Ongur D, Beckmann CF. Using Dual Regression to Investigate Network Shape and Amplitude in Functional Connectivity Analyses. Frontiers in neuroscience. 2017;11:115.

44 Griffanti L, Dipasquale O, Lagana MM, Nemni R, Clerici M, Smith SM, et al. Effective artifact removal in resting state fMRI data improves detection of DMN functional connectivity alteration in Alzheimer’s disease. Front Hum Neurosci. 2015;9:449.

45 Winkler AM, Ridgway GR, Webster MA, Smith SM, Nichols TE. Permutation inference for the general linear model. Neuroimage. 2014;92:381–97.

46 Smith SM, Nichols TE. Threshold-free cluster enhancement: addressing problems of smoothing, threshold dependence and localisation in cluster inference. Neuroimage. 2009;44(1):83–98.

47 Kafkas A, Montaldi D. How do memory systems detect and respond to novelty? Neuroscience Letters. 2018;680:60–68.

48 Marquand AF, De Simoni S, O’Daly OG, Williams SC, Mourao-Miranda J, Mehta MA. Pattern classification of working memory networks reveals differential effects of methylphenidate, atomoxetine, and placebo in healthy volunteers. Neuropsychopharmacology. 2011;36(6):1237–47.

49 Tost H, Braus DF, Hakimi S, Ruf M, Vollmert C, Hohn F, et al. Acute D2 receptor blockade induces rapid, reversible remodeling in human cortical-striatal circuits. Nat Neurosci. 2010;13(8):920–2.

50 Cole DM, Beckmann CF, Oei NY, Both S, van Gerven JM, Rombouts SA. Differential and distributed effects of dopamine neuromodulations on resting-state network connectivity. Neuroimage. 2013;78:59–67.

51 Barkhatova VP. The Role of Catecholamines in the Regulation of Motor Functions. Zh Nevropatol Psikh. 1985;85(7):1068–74.

52 Bart O, Daniel L, Dan O, Bar-Haim Y. Influence of methylphenidate on motor performance and attention in children with developmental coordination disorder and attention deficit hyperactive disorder. Res Dev Disabil. 2013;34(6):1922–7.

53 Auriel E, Hausdorff JM, Herman T, Simon ES, Giladi N. Effects of methylphenidate on cognitive function and gait in patients with Parkinson’s disease: a pilot study. Clin Neuropharmacol. 2006;29(1):15–7.

54 Devilbiss DM, Berridge CW. Low-dose methylphenidate actions on tonic and phasic locus coeruleus discharge. Journal of Pharmacology and Experimental Therapeutics. 2006;319(3):1327–35.

55 Kline RL, Zhang S, Farr OM, Hu S, Zaborszky L, Samanez-Larkin GR, et al. The Effects of Methylphenidate on Resting-State Functional Connectivity of the Basal Nucleus of Meynert, Locus Coeruleus, and Ventral Tegmental Area in Healthy Adults. Frontiers in Human Neuroscience. 2016;10.

56 Engert V, Pruessner JC. Dopaminergic and Noradrenergic Contributions to Functionality in ADHD: The Role of Methylphenidate. Current Neuropharmacology. 2008;6(4):322–28.

57 Xing B, Li YC, Gao WJ. Norepinephrine versus dopamine and their interaction in modulating synaptic function in the prefrontal cortex. Brain Research. 2016;1641:217–33.

58 Bymaster FP, Katner JS, Nelson DL, Hemrick-Luecke SK, Threlkeld PG, Heiligenstein JH, et al. Atomoxetine increases extracellular levels of norepinephrine and dopamine in prefrontal cortex of rat: A potential mechanism for efficacy in Attention Deficit/Hyperactivity Disorder. Neuropsychopharmacology. 2002;27(5):699–711.

59 Hernaus D, Mehta MA. Prefrontal cortex dopamine release measured in vivo with positron emission tomography: Implications for the stimulant paradigm. Neuroimage. 2016;142:663–67.

60 Ye R, Rua C, O’Callaghan C, Jones PS, Hezemans F, Kaalund SS, et al. An in vivo Probabilistic Atlas of the Human Locus Coeruleus at Ultra-high Field. bioRxiv. 2020: 2020.02.03.932087.

61 Glimcher PW. Understanding dopamine and reinforcement learning: The dopamine reward prediction error hypothesis (vol 108, pg 15647, 2011). Proceedings of the National Academy of Sciences of the United States of America. 2011;108(42):17568–68.

62 Torres GE, Gainetdinov RR, Caron MG. Plasma membrane monoamine transporters: Structure, regulation and function. Nature Reviews Neuroscience. 2003;4(1):13–25.

63 Giros B, Wang YM, Suter S, Mcleskey SB, Pifl C, Caron MG. Delineation of Discrete Domains for Substrate, Cocaine, and Tricyclic Antidepressant Interactions Using Chimeric Dopamine-Norepinephrine Transporters. Journal of Biological Chemistry. 1994;269(23):15985–88.

64 Moron JA, Brockington A, Wise RA, Rocha BA, Hope BT. Dopamine uptake through the norepinephrine transporter in brain regions with low levels of the dopamine transporter: Evidence from knock-out mouse lines. Journal of Neuroscience. 2002;22(2):389–95.

65 Eshleman AJ, Carmolli M, Cumbay M, Martens CR, Neve KA, Janowsky A. Characteristics of drug interactions with recombinant biogenic amine transporters expressed in the same cell type. Journal of Pharmacology and Experimental Therapeutics. 1999;289(2):877–85.

66 De Martino B, Strange BA, Dolan RJ. Noradrenergic neuromodulation of human attention for emotional and neutral stimuli. Psychopharmacology. 2008;197(1):127–36.

67 Ulke C, Rullmann M, Huang J, Luthardt J, Becker GA, Patt M, et al. Adult attention-deficit/hyperactivity disorder is associated with reduced norepinephrine transporter availability in right attention networks: a (S,S)-O-[(11)C]methylreboxetine positron emission tomography study. Transl Psychiatry. 2019;9(1):301.

68 Bostan AC, Dum RP, Strick PL. The basal ganglia communicate with the cerebellum. Proc Natl Acad Sci U S A. 2010;107(18):8452–6.

69 Hurley MJ, Mash DC, Jenner P. Markers for dopaminergic neurotransmission in the cerebellum in normal individuals and patients with Parkinson’s disease examined by RT-PCR. European Journal of Neuroscience. 2003;18(9):2668–72.

70 Naatanen R, Michie PT. Early Selective-Attention Effects on the Evoked-Potential - a Critical-Review and Reinterpretation. Biological Psychology. 1979;8(2):81–136.

71 Restuccia D, Della Marca G, Valeriani M, Leggio MG, Molinari M. Cerebellar damage impairs detection of somatosensory input changes. A somatosensory mismatch-negativity study. Brain : a journal of neurology. 2007;130:276–87.

72 Ito M. Opinion - Control of mental activities by internal models in the cerebellum. Nature Reviews Neuroscience. 2008;9(4):304–13.

73 Dominguez-Borras J, Trautmann SA, Erhard P, Fehr T, Herrmann M, Escera C. Emotional Context Enhances Auditory Novelty Processing in Superior Temporal Gyrus. Cerebral Cortex. 2009;19(7):1521–29.

74 Hunkin NM, Mayes AR, Gregory LJ, Nicholas AK, Nunn JA, Brammer MJ, et al. Novelty-related activation within the medial temporal lobes. Neuropsychologia. 2002;40(8):1456–64.

75 Hawco C, Lepage M. Overlapping patterns of neural activity for different forms of novelty in fMRI. Frontiers in Human Neuroscience. 2014;8.

76 Valentini V, Cacciapaglia F, Frau R, Di Chiara G. Differential alpha-mediated inhibition of dopamine and noradrenaline release in the parietal and occipital cortex following noradrenaline transporter blockade. J Neurochem. 2006;98(1):113–21.

77 Valentini V, Frau R, Di Chiara G. Noradrenaline transporter blockers raise extracellular dopamine in medial prefrontal but not parietal and occipital cortex: differences with mianserin and clozapine. Journal of Neurochemistry. 2004;88(4):917–27.

78 van den Brink RL, Pfeffer T, Warren CM, Murphy PR, Tona KD, van der Wee NJA, et al. Catecholaminergic Neuromodulation Shapes Intrinsic MRI Functional Connectivity in the Human Brain. Journal of Neuroscience. 2016;36(30):7865–76.

79 Hernaus D, Marta M, Santa C, Offermann JS, Van Amelsvoort T. Noradrenaline transporter blockade increases fronto-parietal functional connectivity relevant for working memory. European Neuropsychopharmacology. 2017;27(4):399–410.

80 Robison LS, Ananth M, Hadjiargyrou M, Komatsu DE, Thanos PK. Chronic oral methylphenidate treatment reversibly increases striatal dopamine transporter and dopamine type 1 receptor binding in rats. Journal of Neural Transmission. 2017;124(5):655–67.

81 Fusar-Poli P, Rubia K, Rossi G, Sartori G, Balottin U. Striatal Dopamine Transporter Alterations in ADHD: Pathophysiology or Adaptation to Psychostimulants? A Meta-Analysis. American Journal of Psychiatry. 2012;169(3):264–72.

82 Chen W, Liu P, Volkow ND, Pan Y, Du C. Cocaine attenuates blood flow but not neuronal responses to stimulation while preserving neurovascular coupling for resting brain activity. Molecular Psychiatry. 2016;21(10):1408–16.

83 Sigurdardottir HL, Kranz GS, Rami-Mark C, James GM, Vanicek T, Gryglewski G, et al. Association of norepinephrine transporter methylation with in vivo NET expression and hyperactivity-impulsivity symptoms in ADHD measured with PET. Mol Psychiatry. 2019.

